# Individuality, as well as genotype, affects characteristics and temporal consistency of courtship songs in male mice

**DOI:** 10.1101/2021.01.19.427240

**Authors:** Luca Melotti, Sophie Siestrup, Maja Peng, Valerio Vitali, Daniel Dowling, Norbert Sachser, Sylvia Kaiser, S. Helene Richter

**Affiliations:** Department of Behavioural Biology, University of Münster, Badestrasse 13, 48149 - Münster, Germany; Institute for Evolution and Biodiversity, University of Münster, Hüfferstrasse 1, 48149 – Münster, Germany

**Keywords:** Laboratory Mouse, Genetic Background, Strain Differences, Ultrasonic Vocalisations, Intraindividual Variability, Personality, Signature Calls, Social Communication, Mating Strategy, Song Syntax

## Abstract

Courtship songs in mice have been investigated to understand the mechanisms and ecological relevance of vocal communication. There is evidence that courtship song characteristics vary between different genotypes, but little is known on whether individuals, even within the same genotype, differ from each other in the composition, complexity, and temporal consistency of their songs. In a first study, we aimed to systematically identify song features typical of different genotypes, by assessing the composition and complexity (i.e., entropy) of the syllabic sequences of male laboratory mice from four different strains *(Mus musculus* f. *domestica:* C57BL/6J, BALB/c, DBA/2 and B6D2F1). Mice were individually presented with a swab containing fresh female urine for 5 minutes to elicit courtship songs. The four strains differed not only in the composition but also in the complexity of their syllabic sequences. In a second study, we investigated within-strain individual differences in temporal consistency and recurring motifs (i.e., identical sets of syllables that are repeated within a song), using BALB/c and DBA/2 mice. The same procedure as in the first study was followed, but in addition testing was repeated weekly over three weeks. Both strains showed some level of individual temporal consistency; BALB/c in the overall amount of emitted vocalisations and DBA/2 in the expression of specific syllable types. However, hierarchical cluster analysis revealed remarkable individual variability in how consistent song characteristics were over time. Furthermore, recurring motifs were expressed at varying levels depending on the individual. Taken together, not only genotype but also individuality can affect variability in courtship songs in mice, suggesting the existence of different courtship strategies (e.g., higher song consistency to facilitate individual identification) related to varying levels of behavioural plasticity.

**HIGHLIGHTS:**

- Courtship songs in mice can serve as a model to study vocal communication
- We explore how genotype and individuality affect courtship songs’ characteristics
- Genotypes differ in composition and also in complexity of syllabic sequences
- We find remarkable individual variability in how consistent songs are over time
- Results suggest the existence of variation in male courting behaviour

## INTRODUCTION

Auditory signals are one of the most universal and important means of communication across animal species (Portfors & Perkel, 2014). One form, vocal communication, is defined by specific frequency and temporal properties of the vocal signals. Within the context of courtship and mating, some species produce long sequences of vocalisations, also termed “songs” (Holy & Guo, 2005). Songs emitted for courtship serve the function of signalling the presence and identity of the courting individual, in order to attract a potential mate (Simmons et al., 2002).

Mice *(Mus musculus* f. *domestica)* emit vocalizations in a variety of social contexts, including courtship (Ehret, 2018; Portfors, 2007). In their seminal study, Holy and Guo (2005) demonstrated that the ultrasonic vocalisations of male mice during courtship have the characteristics of a song similar to those found in birds or insects but in only few species of mammals (i.e., humans, whales and bats). These songs consist of different syllable types, organised in repeated phrases (i.e., sequences of syllables emitted in close succession; Holy & Guo, 2005). Courtship songs in male mice have been studied to give insight into the ecological relevance of such behaviour and its relationship to reproductive success (e.g., Chabout et al., 2015; Musolf et al., 2010; Nicolakis et al., 2020). Yet the ecological relevance of courtship songs in laboratory mice has been questioned by some authors (Musolf et al., 2010).

The genome of the most common mouse laboratory strains mainly derives from the *Mus musculus domesticus* subspecies, with minor contributions from other subspecies, including *Mus musculus musculus* (Yang et al., 2007, 2011). Laboratory strains were subjected to substantial inbreeding at various stages (Chesler, 2014; Yang et al., 2011), thus reducing the overall genetic and phenotypic variation compared to wild subspecies. The courtship vocalisation repertoire of different laboratory strains, together with the one of wild-derived mice, has only been investigated more in depth in recent years (e.g., Hoffmann et al., 2012; Sugimoto et al., 2011; van Segbroeck et al., 2017), and it is still unclear how exactly the process of artificial selection has affected it. So far, studies conducted with laboratory mice have indicated that male courtship songs emitted in response to fresh female urine have greater syntactical complexity compared to songs produced in the direct presence of a female, and that females are more attracted by and show preference for the urine-related, more complex songs (Chabout et al., 2015). A first study comparing the courtship songs of several laboratory and wild-derived male mice in response to the direct presence of a female indicated that certain syllabic features and their rhythm reduce rejection behaviour by the female and are preferred by females when played back to them (Sugimoto et al., 2011). However, we do not know whether the same syllabic features would also be effective in attracting a female from distance (i.e. via songs triggered by fresh female urine) or whether there are other syntactical characteristics that could influence female preference. Thus, further research is needed to systematically identify the song features typical of a strain and/or individual.

A very small number of studies indicate that courtship songs may have specific characteristics unique to individuals, even when these individuals have the same genotype. The concept of individuality applied to courtship songs can be regarded as the propensity of an individual to maintain similar song characteristics over time and / or across different contexts, (cf. to broader definitions of animal personality; Carere & Maestripieri, 2013). The number of phrases emitted in courtship songs by wild-derived mice has been found to correlate across different social contexts, including the encounter between male and female from either the same or a different population (von Merten et al., 2014). Further, some quantitative features of simpler syllable types (i.e., those syllables without steps), such as duration and average pitch, were suggested to be good indicators of individual identity in courtship songs induced by fresh female urine (Hoffmann et al., 2012). However, the results of this study were based on only one 90 minute session, thus the consistency over time of these individual characteristics was not tested. Only one study so far has shown some level of individual consistency across repeated test sessions for the syllabic composition and the transition probabilities within syllabic sequences (Holy & Guo, 2005), but these findings were based on a small sub-sample of animals and test trials. Taken together, there is a considerable lack of knowledge on the influence of individuality on courtship songs in mice. Concerning the adaptive value, individuality in courtship songs may play an important role in reflecting different courting strategies towards reproductive success (e.g., higher consistency of song characteristics to facilitate individual identification and choice by the female). Thus, if individuality explains at least part of the variation in courtship songs, it is important to assess to what extent this is the case, and to identify those song characteristics that are unique to the individual.

Using a syntactical analysis approach (e.g., Chabout et al., 2015), the aim of this study was twofold. First, we aimed at adding confirmatory evidence of the effects of the genetic background of adult male mice on courtship songs. To do this, we presented males from four different laboratory strains (C57BL/6J, BALB/c, DBA/2 and B6D2F1) with fresh female urine and assessed not only the syllabic composition of the resulting song sequences, but also the complexity of these sequences, using more novel techniques (i.e., entropy analysis; Experiment 1). Second, we selected two strains (BALB/c and DBA/2) based on the information obtained from Experiment 1, and we investigated whether mice having the same genotype showed individual differences in how consistent their syllabic sequences were over time, and in their expression of recurring motifs (i.e., identical sets of syllables that are repeated within a song; Experiment 2). Based on the limited previous evidence, we expected individuality to emerge even within the same genotype.

## METHODS

### Animal Housing and Husbandry

The subject male mice from both experiments (Experiment 1: N = 24; Experiment 2: N = 24; see Experimental Design section for details on strain composition) were purchased from Charles River Laboratories, Sulzfeld, Germany and were delivered to the Department of Behavioural Biology, University of Münster, Germany, at postnatal day (PND) 28. Until PND 63 they were housed in groups of three animals from the same strain, then they were individually housed in transparent Makrolon type III cages (l × b × h: 37 cm × 21 cm × 15 cm) until the end of the experiment. Mice were tested between PND 132 – 232, after undergoing a battery of non-invasive behavioural phenotyping tests as part of a separate study (PND 76 – 93).

The female mice used for the social encounter sessions and for urine collection originated from the stock of the Department of Behavioural Biology, University of Münster, Germany (5- HTT +/− and 5-HTT +/+ mice with a C57BL/6J genetic background; Experiments 1 and 2) or were purchased at PND 21 from Charles River Laboratories, Sulzfeld, Germany (Experiment 2; see appendix A1, Social Encounter and Urine Collection sub-sections). They were housed in groups of two to five females in transparent Makrolon type III cages with enrichment (see appendix A1).

All animals had *ad libitum* access to water and food pellets (Altromin 1324, Altromin Spezialfutter GmbH & Co. KG, Lage, Germany) and were kept at a room temperature of about 22°C and relative humidity of about 50%. For details of cage furnishing and enrichment see appendix A1. A 12:12 light:dark cycle (lights off at 9:00 a.m. for males and at 10:00 a.m. for females) was maintained and experimental procedures were conducted during the dark phase under red light. Male and female mice were housed in separate rooms.

All procedures complied with the regulations covering animal experimentation within Germany (Animal Welfare Act) and the EU (European Communities Council DIRECTIVE 2010/63/EU) and were approved by the local (Amt für Gesundheit, Veterinär- und Lebensmittelangelegenheiten, Münster, Nordrhein-Westfalen, reference number: 39.32.7.1) and federal authorities (Landesamt für Natur, Umwelt und Verbraucherschutz Nordrhein- Westfalen “LANUV NRW”).

Individual ear cuts were used to identify all mice. Each mouse received one partial punch at the margin of one ear (no risk of injury / getting stuck) after mild anaesthesia using specific ear punch forceps. No swelling or bleeding was noticed. At the end of the experiment, mice remained within our facility or were handed over to a cooperation partner.

### Experimental Design

This study consisted of two experiments. Experiment 1 aimed at assessing whether the genetic background of male mice from four different strains affected the syllabic composition and complexity of courtship songs. Experiment 2 investigated individuality in the consistency over time of song characteristics and in the expression of recurring motifs in two different mouse strains.

#### Experiment 1

Subjects were 24 adult male mice from four different strains (six mice per strain), namely the inbred strains C57BL/6J, BALB/c and DBA/2 and the hybrid strain B6D2F1 (C57BL/6J × DBA/2). The experiment was carried out in three partially overlapping batches, where two male mice of each strain were randomly assigned to each batch (Fig. 1). Between PND 132 – 136, each mouse experienced a 20 minute social encounter with an adult female mouse (genetic background: C57BL/6J), which has been shown to elicit the emission of courtship ultrasonic vocalisations in males (Dizinno et al., 1978; Nyby et al., 1978; Sipos et al., 1992). One week later, testing was conducted over two consecutive days and consisted of one test session and one control session, the order of which was counterbalanced for each strain and batch. During the test session mice were individually presented with fresh female urine on a cotton swab for five minutes in order to elicit the emission of courtship songs; in the control session they were presented with a clean cotton swab without urine to control for any effect of the experimental procedure on vocalisations emission. Vocalisations emitted during both sessions were recorded and analysed. For further details on vocalisations recording setup and experimental procedures (social encounter, test / control sessions, and urine collection) see appendix A1.

**Figure 1.**
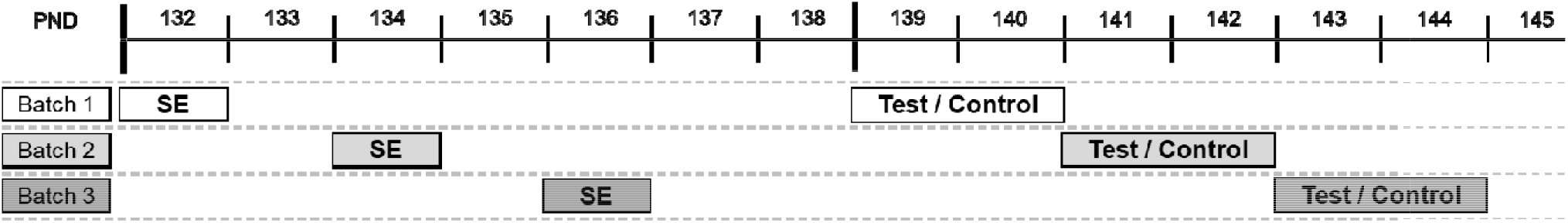
Timeline of Experiment 1. The experiment was carried out in three partially overlapping batches during a time period of two weeks. Each batch was comprised of two male mice from each of the C57BL/6J, BALB/c, DBA/2 and B6D2F1 strains. All males experienced a 20 minute social encounter (SE) with an adult female with a C57BL/6J genetic background. Seven days later, males were individually presented with either a cotton swab containing fresh female urine (test session) or a clean cotton swab without urine (control session). Control and test sessions were performed over two consecutive days, and their order was counterbalanced within each strain.

#### Experiment 2

Subjects were 24 adult male mice from the BALB/c and DBA/2 strains (12 mice per strain). These two strains were chosen based on the differences in syllabic composition found in Experiment 1 and on the relatively high number of vocalisations emitted. The experimental procedures were carried out between PND 188 – 231 and were the same as in Experiment 1, but this time males underwent three weekly test sessions (instead of only one test session) to assess individual consistency over time in courtship song features. Also, the control session always preceded the first test session, based on evidence from Experiment 1 of a carry-over effect in the transition between first test session and control session (see appendix A1, Comparison of Control and Test Sessions). The experiment was carried out in four partially overlapping batches, where three mice of each strain were randomly assigned to each batch (Fig. 2; see appendix A1, Experimental Procedures section, for more details).

**Figure 2.**
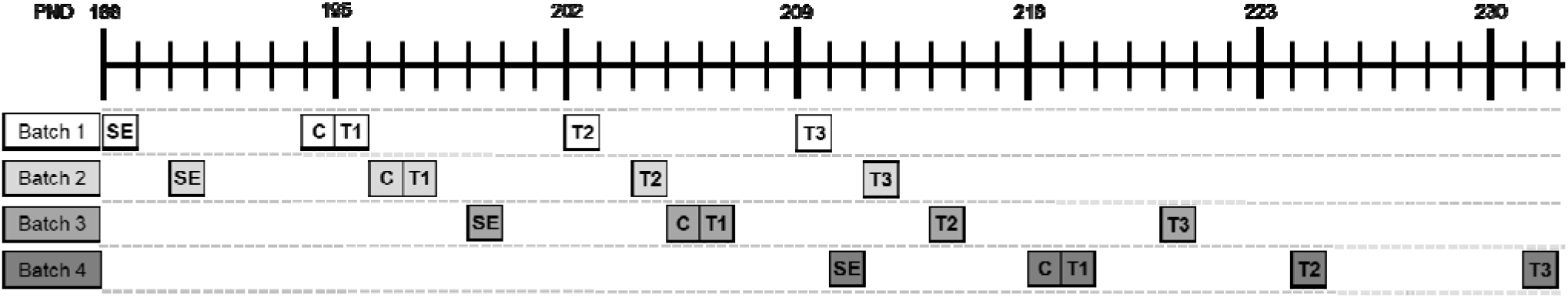
Timeline of Experiment 2. This experiment was carried out in four partially overlapping batches, where three adult male mice of two strains (BALB/c and DBA/2) were randomly assigned to each batch. Every mouse experienced a 20 minute social encounter (SE) with an adult female with a C57BL/6J genetic background. Six days later, a control session (C) was performed where the animal was presented with a clean cotton swab without urine for five minutes. On the following day and weekly for three weeks (T1, T2 and T3), each mouse was presented with fresh female urine on a cotton swab for five minutes in order to elicit the emission of courtship songs.

### Vocalisations Analysis

All vocalisation recordings were blinded and their order randomised to avoid experimenter bias during the analysis. Vocalisations were analysed using the software Avisoft – SASLab Pro (Version 5.2.10; Avisoft Bioacoustics, Germany). Spectrograms were generated with a FFT-length of 512 points (e.g., Musolf et al., 2010) and a time window overlap of 75 % (e.g., Scattoni et al., 2008). The frame size was 100 % and a flat-top window was used (e.g., Musolf et al., 2010).

In Experiment 1, the whole five minutes recorded for each test / control session were analysed. Based on the finding in Experiment 1 that BALB/c and DBA/2 mice emitted relatively high levels of vocalisations, in Experiment 2 only the last four minutes (out of the five minutes recorded) of each session were analysed. We excluded the first minute based on the observation that not all animals started to vocalise at the very start of the recording session.

Based on previous literature (Grimsley et al., 2011), 12 syllables types were identified and counted manually using the “standard marker” (for durations) and “free reticular” (for frequencies) cursors within the Avisoft – SASLab Pro software. Definitions of the syllable types and examples of their spectrograms are provided in Table 1 and Fig. 3, respectively. The total number of emitted syllables and the percentages of syllable types on the total number of syllables were calculated for each recording session. Furthermore, the sequences of emitted syllables within each recording session were recorded in temporal order to allow for sequence analysis.

**Table 1.**
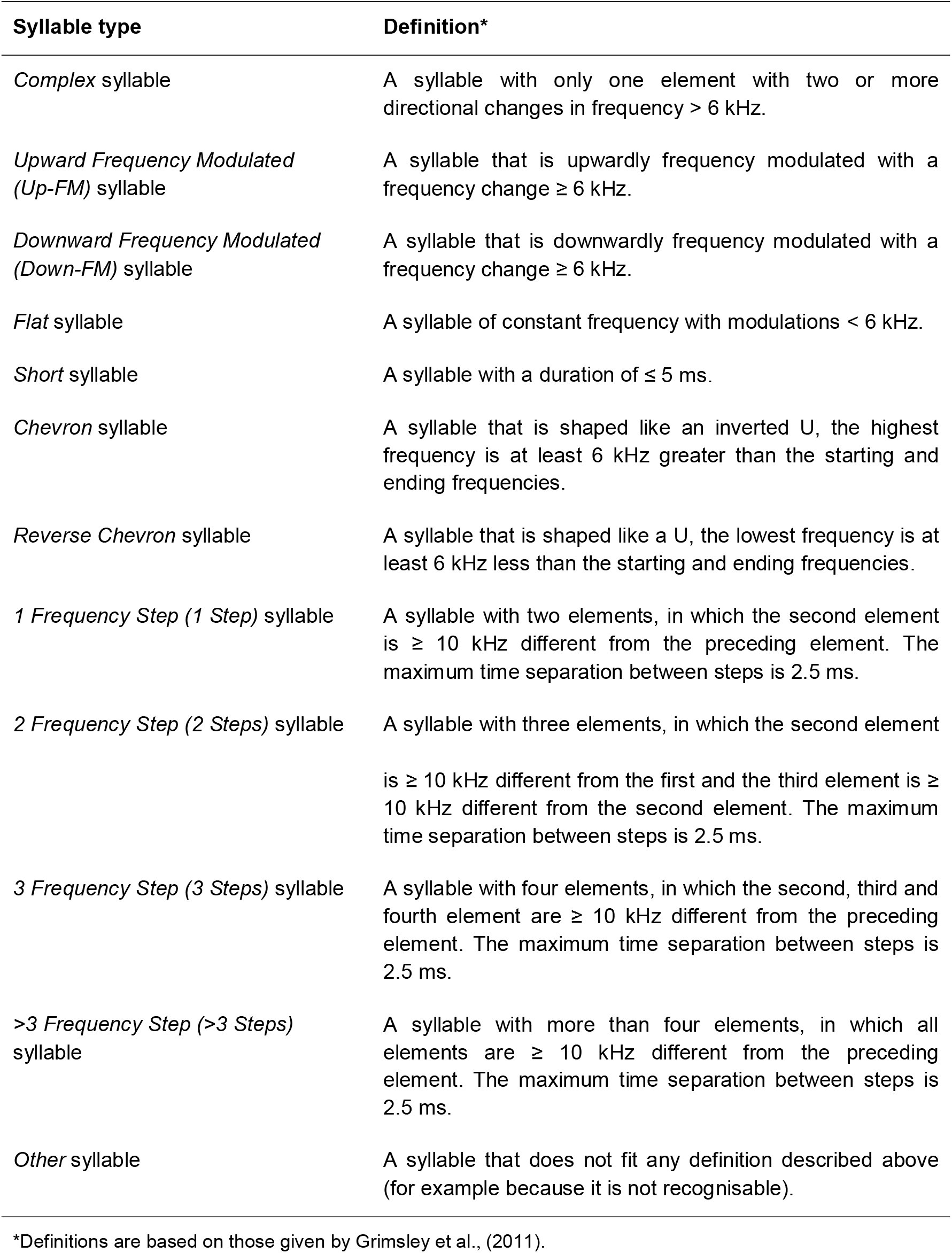
Definitions of counted syllable types.

**Figure 3.**
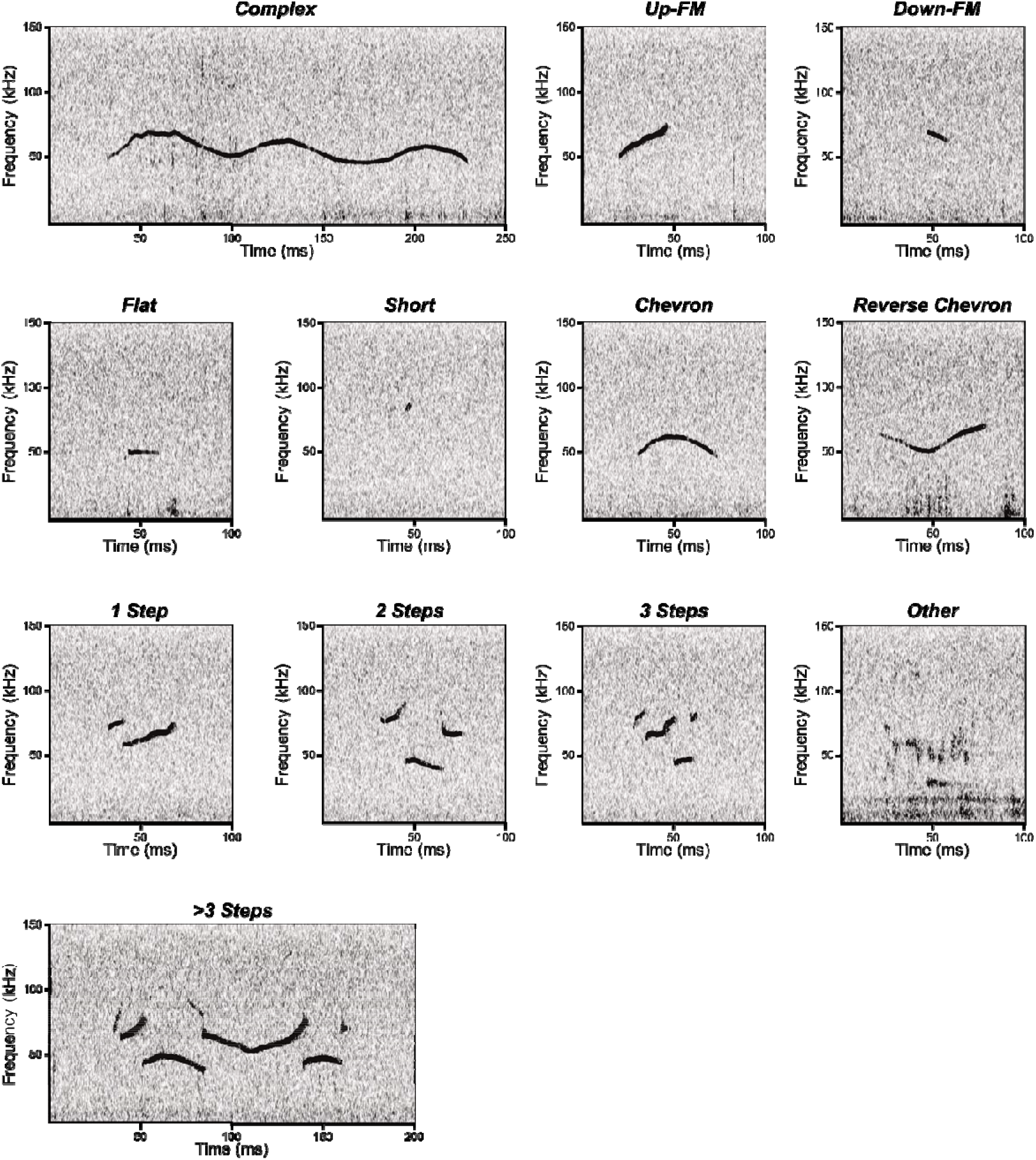
Spectrogram examples of each syllable type identified and counted during the analysis of the recordings. The x axis indicates the duration of the syllable and the y axis its frequency (kHz).

### Data Analysis

Data were analysed using IBM SPSS (version 25) and R (version 3.4.4). In Experiment 1, the comparisons between control and test sessions and the analysis of strain differences in syllabic composition and sequence complexity were carried out using non-parametric tests, as the data did not meet the assumptions for parametric statistics. To allow for better comparability of results between Experiments, the same was done for Experiment 2. Data for these variables were thus summarised as median (M) and interquartile range (IQR). Principal component analysis was used to provide a visual summary of the strain differences in syllabic composition (Experiment 1). Consistency of song characteristics across time was assessed by calculating intraclass correlation coefficients; similarity between song sequences, within both strain and individual, was visualised using hierarchical cluster analysis (Experiment 2). See below for a detailed description of the data analysis.

### Measures of Internal Validity

Intra- and inter- rater reliability of syllable counts was assessed using intraclass correlation coefficients (see appendix A1 for details).

To confirm that courtship songs were emitted specifically in the presence of female urinary cues, control and test sessions were compared using related-samples Wilcoxon signed rank tests (Experiment 1) and related-samples Friedman’s tests (Experiment 2; see appendix A1 for details).

### Strain Differences in Syllabic Composition and Sequence Complexity

In Experiment 1, independent samples Kruskal-Wallis tests were used to compare the total numbers of syllables and the percentages of different syllable types emitted during the test sessions across the four strains. Because of the positive dependency of the p values obtained with the Kruskal-Wallis tests, the p values were corrected with the Benjamini-Hochberg procedure, allowing a false discovery rate of 10 % (Benjamini & Hochberg, 1995; Benjamini & Yekutieli, 2001). In the cases where the Kruskal-Wallis test indicated a significant strain effect, pairwise strain comparisons were carried out via Dunn’s post hoc tests, and the resulting p values were adjusted using the Bonferroni correction for multiple comparisons (Armstrong, 2014). Strain differences were visualised using principal component analysis (PCA), which was performed with the R package *FactoMineR* (version 2.3). The relative frequencies of syllable types from each song were used to construct the PCA object. Values were scaled to unit variance before multidimensionality reduction. The PCA output was visualised with the R package *factoextra* (version 1.0.7).

In Experiment 2, differences in total numbers of syllables and in percentages of different syllable types (average across the three test sessions) between BALB/c and DBA/2 were assessed using Mann-Whitney U tests. P values were corrected with the Benjamini-Hochberg procedure as described for Experiment 1.

Complexity of the song sequences was investigated in both experiments. A Shannon entropy analysis *(DescTools* package in R) was performed and entropy scores were generated, with and without correction for sequence length. Correction for sequence length consisted of dividing the entropy score by the total number of syllables emitted within the same song. Relatively higher entropy scores indicated greater complexity (e.g., greater variability in syllable expression) of the song sequence. The effect of strain on sequence complexity was assessed using an independent samples Kruskal-Wallis test (four strains, Experiment 1) and a Mann-Whitney U test (two strains, Experiment 2). In Experiment 2, the entropy scores from the three test sessions were averaged.

### Individuality in Consistency over Time, Sequence Complexity and Recurring Motifs

Experiment 2, being comprised of three repeated test sessions, allowed for the investigation of individual differences in temporal consistency, complexity (entropy) and recurring motifs of the song sequences.

Individual consistency over time in total number of syllables and in percentages of syllable types across the three test sessions was assessed using the intraclass correlation coefficient (ICC). For each strain, the total number of syllables and the four most frequent syllable types were analysed. The ICC was calculated using a two way mixed design assessing the consistency of the mean of the test sessions (Koo & Li, 2016; Landers, 2015; Shrout & Fleiss, 1979). Consistency was evaluated based on the ICC estimate and on the lower bound of the 95% confidence interval (Koo & Li, 2016). To graphically show individuality based on similarity between song sequences, hierarchical cluster analysis was used. The distances between song sequences were calculated using the *stringdist* package in R using the qgram method (q = 3 consecutive syllables). This method uses sub-sequences (qgrams) of a defined length (q) to calculate the similarity between two or more sequences. The distances between each song sequence were then used to build a dendrogram.

To assess individual consistency in complexity of the song sequences, intraclass correlation coefficients (ICC) were calculated for each strain using the individual entropy scores from the three test sessions. ICC calculation and interpretation were the same as for the assessment of consistency in syllabic composition.

In order to investigate individual variation in the production of recurring motifs within courtship songs, we implemented an *ad hoc* algorithm in R (see S1). This algorithm extracted all the identical sub-strings (motifs) that had a minimum length of four syllables and that were repeated at least four times within the same song, and tested their statistical overrepresentation. Briefly, the algorithm reshuffled each song sequence multiple times (n = 1000 randomisation replicates) and counted the occurrence of motifs in the reshuffled sequences. For each motif, the total motif count across randomisation replicates and its sample space were used to calculate the expected frequency of motifs in the randomised sequences. The sample space (number of all possible sequence substrings of a given length) was calculated as: (sequence length – motif length) * randomization replicates. The statistical overrepresentation of the observed motifs was computed with one-sided binomial tests, so that only the motifs that occurred significantly more often than chance were retained. p values were adjusted for multiple comparisons using Bonferroni correction, which was performed on a *per*-sequence basis, with the number of comparisons equal to the number of motifs observed in each song sequence. This motif extraction procedure led to a final sample of 337 selected recurring motifs across all song recordings. The number of unique motifs produced *per* animal and test session was counted as a measure of individual variability in motif expression. Furthermore, the ratio between the number of unique syllables in each motif and the total number of syllables in the same motif was calculated as a measure of motif complexity. Ratios that belonged to the same animal were averaged to obtain a motif complexity score *per* animal. Finally, the number of repetitions of each motif within a test session was counted. Counts that belonged to the same animal were averaged to obtain a motif repetition score *per* animal.

### Exclusion Criteria

In Experiment 1, two out of six C57BL/6J males only emitted four syllables during the test session and were thus excluded from the analysis of syllabic composition.

As for the entropy, principal component, hierarchical cluster, and recurring motifs analyses, the minimum length below which a song sequence was excluded from the analysis was set to 150 syllables, leading to the exclusion of two C57BL/6J mice (Experiment 1) and of four song sequences of BALB/c mice in the third test session (Experiment 2). This threshold was chosen being the shortest substring of a scrambled sequence (random string) for which the syllable frequency distribution is representative of the distribution of the whole sequence. The same criterion was applied for the ICC analysis of syllabic composition (Experiment 2); the fact that only the total number of syllables and the four most frequent syllable types were analysed allowed to set a lower minimum threshold of 50 syllables. This led to the exclusion of two BALB/c mice which emitted less than 50 syllables in the third test session.

## RESULTS

### Strain Differences in Syllabic Composition and Complexity of Song Sequences

#### Experiment 1 – Differences between Four Strains

The total number of syllables emitted during the test session was affected by strain (H(3) = 13.81, Benjamini-Hochberg adjusted p = 0.007), with C57BL/6J mice vocalising less than the other strains (significant Bonferroni-corrected comparisons: C57BL/6J < B6D2F1, DBA/2; trend: C57BL/6J < BALB/c; Fig. 4A). Strain significantly affected the percentages of *Flat* syllables (H(3) = 9.21, adj. p = 0.046; significant Bonferroni-corrected comparisons: C57BL/6J > B6D2F1; Fig. 4E), *Short* syllables (H(3) = 14.39, adj. p = 0.006; significant Bonferroni-corrected comparisons: C57BL/6J > BALB/c, B6D2F1; Fig. 4F), *Chevron* syllables (H(3) = 14.88, adj. p = 0.006; significant Bonferroni-corrected comparisons: BALB/c > C57BL/6J, B6D2F1; Fig. 4G), *2 Steps* syllables (H(3) = 14.89, adj. p = 0.006; significant Bonferroni-corrected comparisons: DBA/2 > C57BL/6J, BALB/c; Fig. 4J), *3 Steps* syllables (H(3) = 15.35, adj. p = 0.006; significant Bonferroni-corrected comparisons: B6D2F1 > BALB/c; Fig. 4K) and *>3 Steps* syllables (H(3) = 9.52, adj. p = 0.046; significant Bonferroni-corrected comparisons: C57BL/6J > BALB/c; Fig. 4L). A trend for a strain effect was found for the percentages of *Down-FM* syllables (H(3) = 7.71, adj. p = 0.07). From visual inspection of Fig. 4D, C57BL/6J mice tended to emit a higher percentage of *Down-FM* syllables than the mice of the other strains. Another trend for a strain effect was found on the percentages of *Reverse Chevron* syllables (H(3) = 7.71, adj. p = 0.07). Visual inspection of Fig. 4H indicates that C57BL/6J and B6D2F1 mice showed a higher percentage of *Reverse Chevron* syllables than BALB/c and DBA/2 mice. There was no significant strain effect on the percentages of *Complex* syllables (H(3) = 5.10, adj. p = 0.20), *Up-FM* syllables (H(3) = 3.54, adj. p = 0.34) and *1 Step* syllables (H(3) = 3.20, adj. p = 0.36; Fig. 4B, 4C, 4I). Fig. 4 also shows that the courtship songs of the hybrid strain B6D2F1 were more similar to those of the maternal strain (DBA/2) overall, indicating a dominant, rather than intermediate, inheritance pattern. However, the expression of syllables with frequency jumps varying in complexity (*2, 3,* and *>3 Steps* syllables) suggests the presence of a rather intermediate inheritance, since B6D2F1 mice produced more *3 Steps* syllables, while the (parental) DBA/2 and C57BL/6J strains emitted more *2 Steps* and *>3 Steps* syllables, respectively. A visual display of the overall effect of strain on the syllabic composition is given in Fig. 5, as a result of principal component analysis. From this figure it can be seen that, overall, strain differences are present with some overlap, confirming previous findings. The C57BL/6J strain appears to differ the most from the other strains.

**Figure 4.**
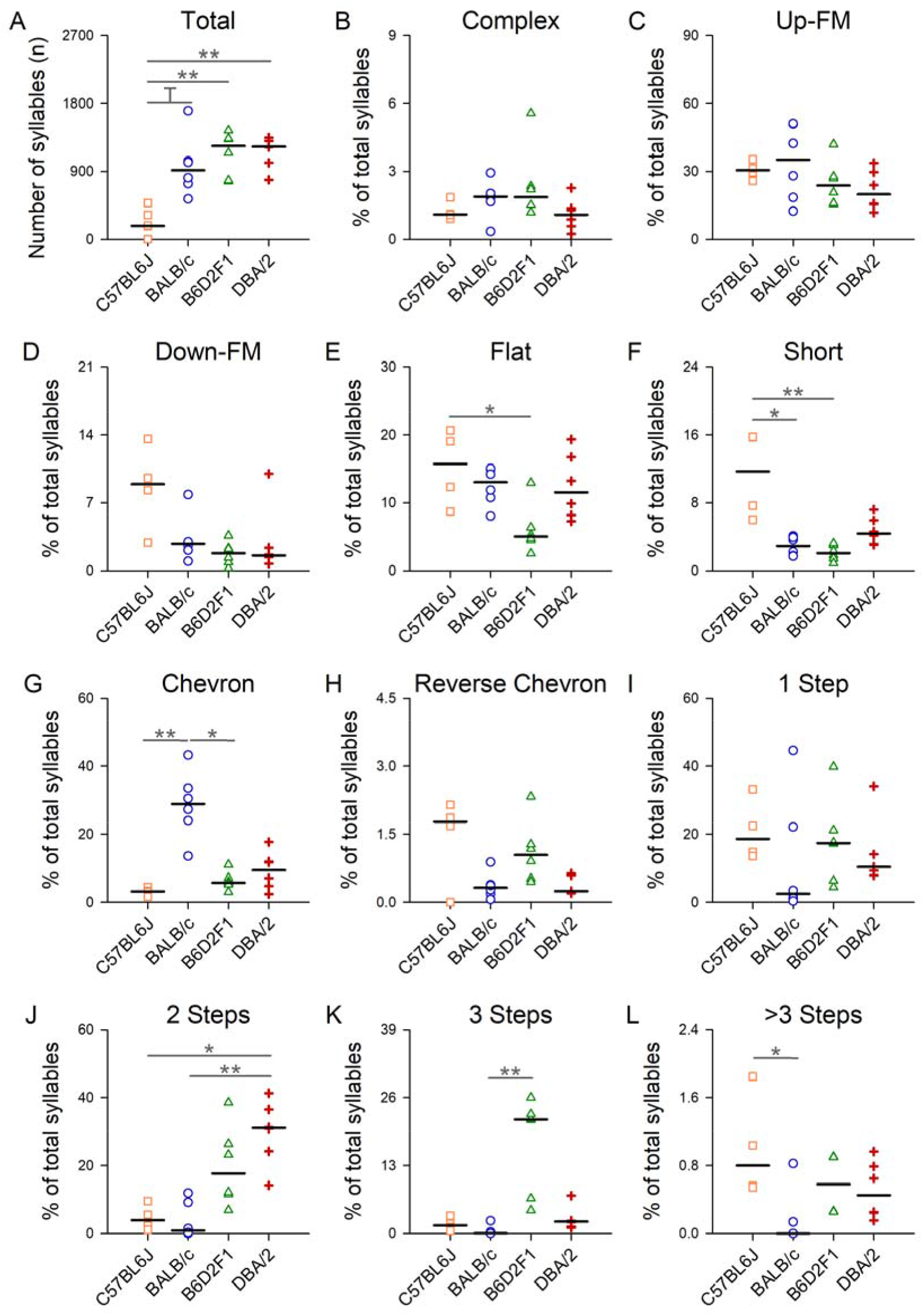
Plots showing strain differences (C57BL/6J, BALB/c, DBA/2 and B6D2F1) in the total number of syllables and in the percentages of different syllable types emitted during the test sessions. Horizontal lines indicate medians, dots show individual data points (N_C57BL/6J_ = 4, N_BALB/C_ = N_B6D2F1_ = N_DBA/2_ = 6). T = p < 0.1, * = p < 0.05, ** = p < 0.01, *** = p < 0.001. Test for strain effect: independent samples Kruskal-Wallis test corrected with Benjamini-Hochberg procedure. Pairwise comparisons: asymptotic Dunn’s post hoc test, p-values adjusted with Bonferroni correction.

**Figure 5.**
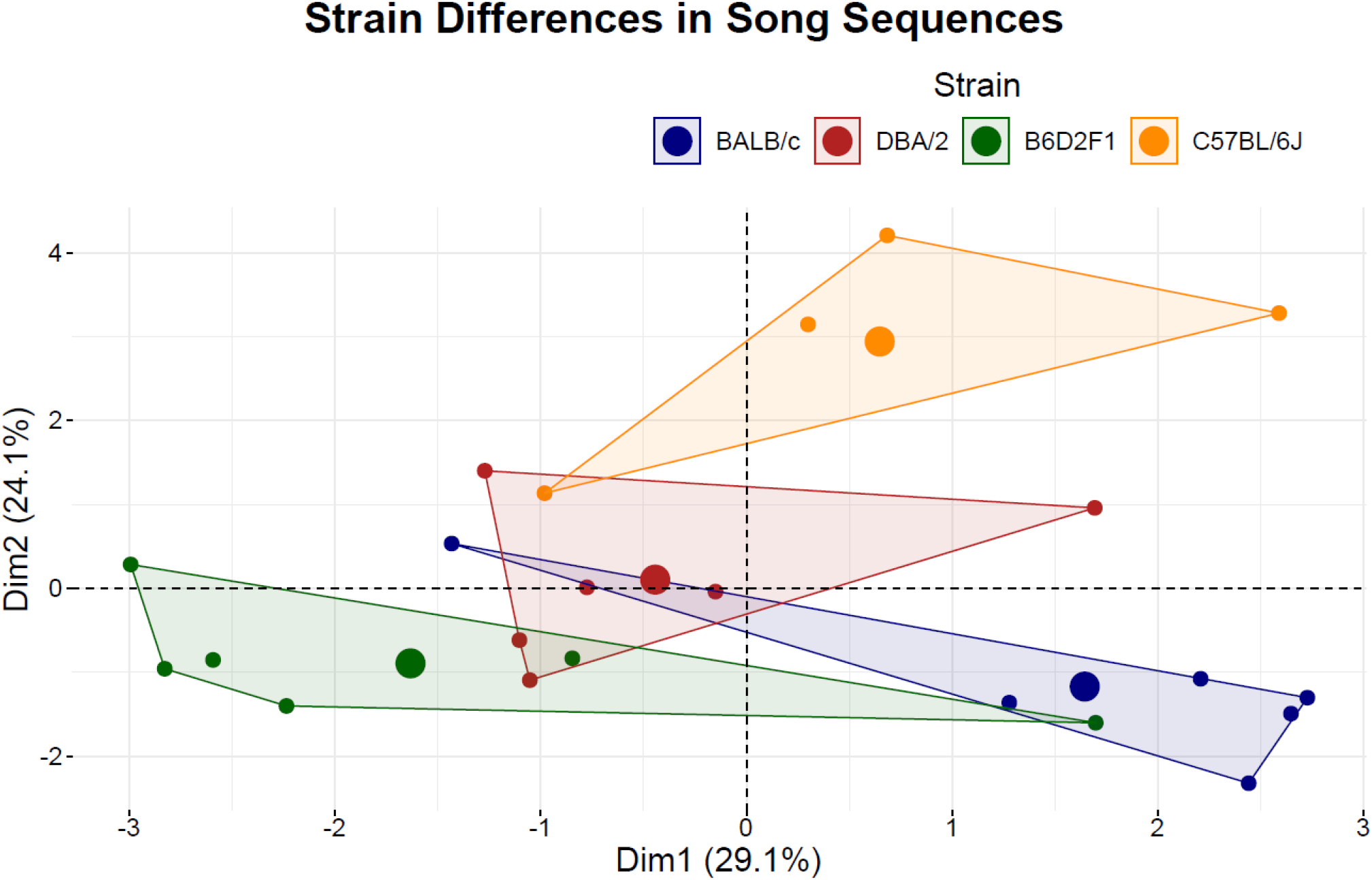
Strain differences in song sequences displayed using principal component analysis of syllable proportions. Animals (individual small dots) from the same strain are assigned the same colour, and the maximum area covered by connecting the individual dots from the same strain is highlighted. Strain differences are shown across the two dimensions explaining the majority of variation (53.3 % of the variance). Larger dots indicate the mean value for each strain. The less the highlighted areas overlap the more different are the strains.

Shannon entropy analysis of song sequences indicated that complexity was lower in the BALB/c strain compared to the other strains (H(3) = 13.00, p = 0.005; significant Bonferroni-corrected comparisons: BALB/c < C57BL/6J, B6D2F1; trend: BALB/c < DBA/2). However, when correcting the entropy scores for song sequence length, it was the C57BL/6J strain which showed relatively higher complexity overall (H(3) = 9.78, p = 0.02; significant Bonferroni-corrected comparisons: C57BL/6J > B6D2F1, DBA/2; appendix A2), suggesting that the extent to which sequence length and sequence entropy contribute to the overall sequence complexity differs between strains.

#### Experiment 2 – Differences between Two Strains

Strain differences between BALB/c and DBA/2 mice in the total number of syllables produced and in the percentages of different syllable types emitted at testing (as average of the three test sessions) are displayed in appendix A3. The same strain differences as in Experiment 1 were observed; in addition, some differences that were only indicated graphically but not statistically in Experiment 1, were found to be statistically significant (with a larger sample size) in Experiment 2.

The total number of syllables emitted during the test sessions was affected by strain (U = 113.0, adj. p = 0.03), with DBA/2 vocalising more than BALB/c overall. BALB/c mice emitted more *Up-FM* (U = 29.0, adj. p = 0.03) and *Chevron* (U = 19.0, adj. p = 0.003) syllable types than DBA/2 mice, while DBA/2 mice produced more *1 Step* (U = 111.0, adj. p = 0.04), *2 Steps* (U = 143.0, adj. p < 0.001), *3 Steps* (U = 142.0, adj. p < 0.001) and *>3 Steps* (U = 141.5, adj. p < 0.001) syllable types than BALB/c. No significant strain differences were found for the syllable types *Complex* (U = 51.5, adj. p = 0.29), *Down-FM* (U = 42.0, adj. p = 0.13), *Flat* (U = 64.0, adj. p = 0.73), *Short* (U = 98.0, adj. p = 0.19) and *Reverse Chevron* (U = 73.0, adj. p = 1.00).

In accordance with Experiment 1, Shannon entropy analysis of song sequences showed that complexity was greater in DBA/2 mice than BALB/c mice (U = 126.0, p = 0.001). However, when correcting the entropy scores for song sequence length, there was no significant difference between the two strains (U = 47.0, p = 0.16), suggesting that sequence length alone explains the differences in sequence complexity between these two strains.

### Individuality in Syllabic Composition, Sequence Complexity and Recurring Motifs

#### Syllabic Composition

In Experiment 2, intraclass correlations revealed that individual temporal consistency of songs’ syllabic composition varied according to the strain. Temporal consistency in BALB/c mice was moderate to excellent for the total number of syllables (ICC_average_ = 0.859; CI lower bound = 0.626), while in DBA/2 mice it was moderate to excellent for the *2 Steps* syllable type (ICC_average_ = 0.823; CI lower bound = 0.531) and good to excellent for the *Up-FM* syllable type (ICC_average_ = 0.915; CI lower bound = 0.775). All the other variables analysed (BALB/c: *Up-FM, Chevron, Flat* and *1 Step* syllable types; DBA/2: Total number of syllables, *Flat* and *1 Step* syllable types) showed poor individual temporal consistency across the three test sessions (all CI lower bounds < 0.5; see appendix A4).

The dendrogram illustrating the similarity between song sequences from the same strain and from the same individual (Fig. 6) illustrates the presence of a gradient across individuals ranging from high to low individual consistency over time in both strains. Examples of high or low temporal consistency in syllabic composition are shown in appendix A5 for individuals of each strain. This suggests a remarkable individual variation in how similar courtship songs are within the same individual, a finding which could be generalised across the two different strains.

**Figure 6.**
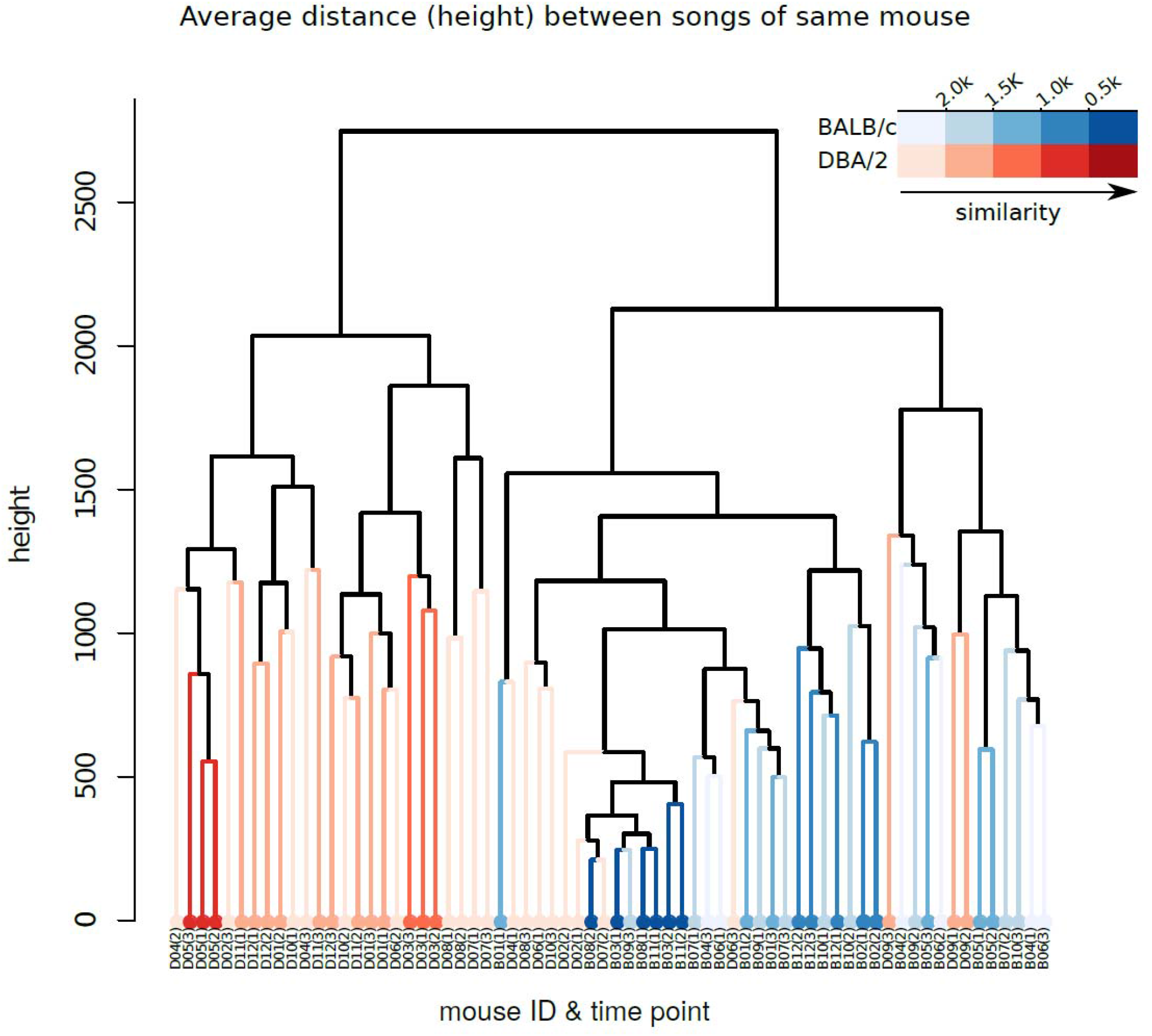
Dendrogram showing the hierarchical clustering of the song sequences from Experiment 2. Each ID corresponds to the song sequence of a mouse during one of the three test sessions. Different colours indicate to which strain the sequence belongs (red: DBA/2; blue: BALB/c), while the colour shade indicates if the sequences are relatively similar (darker shade) or relatively dissimilar (lighter shade) within the same individual. The further away sequences are in the tree (i.e., the higher is the level on the y axis at which two or more sequences are connected), the more dissimilar their syllabic composition is.

#### Sequence Complexity

Intraclass correlation analysis revealed poor individual consistency of entropy scores across the three test sessions in both BALB/c and DBA/2 mice (ICC_average_ = 0.551; CI lower bound = −0.517; ICC_average_ = 0.331; CI lower bound = −0.770; respectively; Fig. 7).

**Figure 7.**
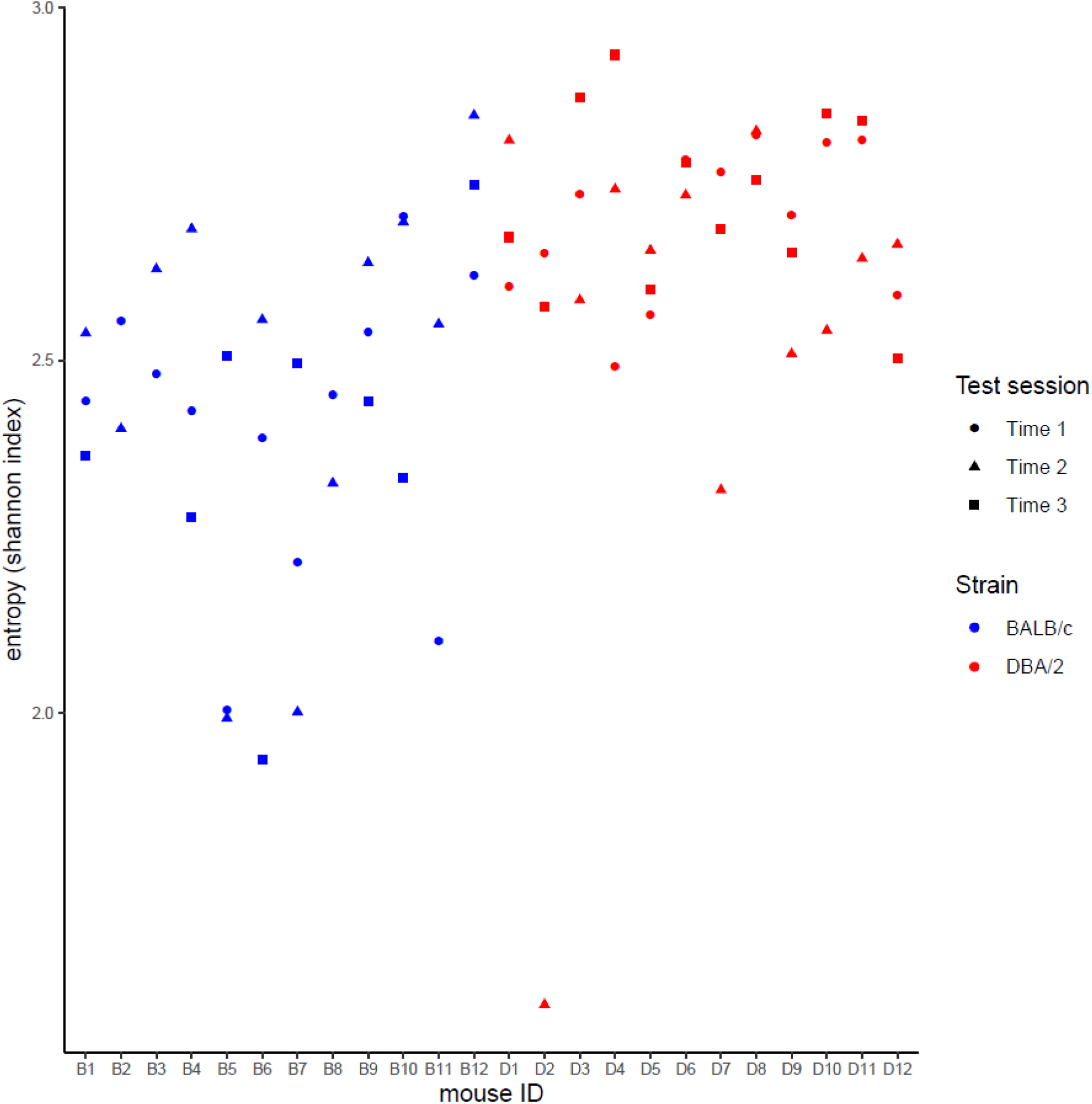
Individual entropy levels across test sessions. Shown are the entropy scores of the three test sessions (dots) for each mouse. Different colours indicate BALB/c and DBA strains, while the different dot shapes specify the test session.

#### Recurring Motifs

The number of unique and non-random recurring motifs identified ranged from 0 to 26 *per* animal and test session, indicating a relatively large variation across animals (mean ± SD; BALB/c: 4.28 ± 4.30; DBA/2: 5.31 ± 2.88). However, the structure of the motifs had low complexity (i.e., only about one third of a motif, on average, comprised unique syllables, while the rest consisted of repetitions of the same syllable) and low variation across animals (BALB/c: 0.344 ± 0.042; DBA/2: 0.367 ± 0.048). Similarly, motif repetition scores indicated low variation across individuals (i.e., each individual repeated the same motif a similar number of times; BALB/c: 5.12 ± 1.04; DBA/2: 4.92 ± 0.54).

## DISCUSSION

This study investigated the influence of genotype and individuality on courtship songs in the male mouse (*Mus musculus* f. *domestica),* with a focus on the syntactical characteristics of the song sequences. The four laboratory strains (C57BL/6J, BALB/c, DBA/2 and B6D2F1) investigated in Experiment 1 differed not only in the composition but also in the complexity (entropy) of their syllabic sequences. In Experiment 2, we found that the syllabic sequences of mice with the same genotype (either BALB/c or DBA/2 strain) showed some level of temporal consistency at the individual level; BALB/c mice in the total number of syllables produced, and DBA/2 mice in the expression of specific syllable types *(2 Steps* and *Up-FM* syllables). However, hierarchical cluster analysis of the syllabic sequences highlighted a remarkable individual variability in consistency across test sessions in both strains. Furthermore, we identified recurring, non-random song motifs which were expressed at different levels depending on the individual; when emitted, motifs were characterised by a simple structure and similar levels of repetition within a song.

### Effect of Genotype on Syllabic Composition and Complexity of Songs

Genotype strongly affected the syllabic composition of courtship songs in both experiments. Our results, consistent with previous literature, indicate that in the laboratory mouse, genotype can be predictive not only of the length of a song sequence (i.e., the rate at which syllables are emitted), but also of which syllables are used and how often. C57BL/6J mice, for instance, produced relatively higher levels of *Short* and *>3 Steps* syllables, in line with findings from previous studies comparing C57BL/6J with BALB/c mice (Kikusui et al., 2011; Sugimoto et al., 2011). However, they vocalised at a lower rate overall, compared to the other strains. Based on principal component analysis, which took into account the variation of the whole syllabic composition of a song, the C57BL/6J strain appeared to differ the most from the other strains in this study. The fact that C57BL/6J mice vocalised the least may indicate that this strain vocalises less in general, as shown in other contexts (male-male interactions: van Segbroeck et al., 2017; mother-pup interactions: Bell et al., 1972; D’Amato et al., 2005). BALB/c mice, instead, emitted relatively higher levels of *Chevron* syllables, supporting previous findings that this specific syllable may be typical of this strain (Kikusui et al., 2011; Sugimoto et al., 2011).

The complexity of the syllabic sequences, measured as sequence entropy, varied across strains and was affected by sequence length (syllable emission rate). C57BL/6J mice emitted syllabic sequences at a lower rate but with higher complexity, while BALB/c mice vocalised at a higher rate but with less complex syllabic sequences, and the DBA/2 and B6D2F1 strains showed intermediate patterns. While the biological significance of this finding should be empirically tested in future studies, our result suggests that both sequence entropy and sequence length should be evaluated together when determining the overall sequence complexity of a genotype.

The observed strain differences in composition and complexity of the syllabic sequences could be used in future studies to better understand how domestication has influenced courtship songs. Only a few studies investigated courtship songs of wild-derived mice (*Mus musculus musculus)* in response to unfamiliar female urine. Comparing these studies with our findings, it appears that wild mice emit courtship vocalisations at a lower rate compared to laboratory strains (all strains combined in our study: M = 916 syllables in 5 min; Musolf et al., 2010: M < 300 syllables in 30 min; Hoffmann et al., 2012: M = 1122 syllables in 90 min). This confirms previous evidence that generally domestication may lower the threshold for the display of courtship behaviour (e.g., Künzl & Sachser, 1999; Price, 1984).

Furthermore, the proportion of syllables which could be classified as “more complex” (i.e., those containing steps; Chabout et al., 2015) appears to be much greater in laboratory strains (Fig. 4 and appendix A3), as only about 2 % of the emitted vocalisations over a 90 minute exposure to unfamiliar female urine consisted of 1 *Step* and 2 *Steps* syllables in wild male mice (Hoffmann et al., 2012). Also, the inter-individual variation in the number of syllables emitted in our laboratory strains appears to be comparable, or only marginally lower, to that of wild-derived mice tested within the same context (see Musolf et al., 2010 for comparison). Therefore, the relatively high complexity and inter-individual variation of courtship songs observed make laboratory mice an ideal model for understanding the basics of vocal communication. In addition, there is some evidence that wild-derived female mice are able to discriminate between courtship songs of different mouse species (Musolf et al., 2015), and that female laboratory mice can distinguish between songs of different strains (Nomoto et al., 2020). Similarly, the typical syllabic composition (and temporal features, Sugimoto et al., 2011) of courtship songs of a laboratory strain, together with the combined assessment of syllabic emission rate and sequence entropy, could be used to assess if females use this information to also discriminate between males from the same strain.

### Individuality in Courtship Songs

When assessing the individual consistency over time of syllabic composition and sequence complexity within the same genotype, we found that the degree of expression of certain syllable types, but not the overall complexity of the song sequence, were repeatable across test sessions. These markers for individuality however differed depending on the strain considered. The consistency over time in the total number of syllables observed in BALB/c mice appears to somehow relate to the finding in wild-derived mice that the number of emitted phrases is consistent across different social contexts (von Merten et al., 2014). Instead, the consistency over time of two specific syllable types (*2 Steps* and *Up-FM* syllables) observed in DBA/2 mice partly supports the preliminary finding by Holy and Guo (2005) that for a sub-sample of B6D2F1 mice, the choice of syllable types was consistent over the same three week period as in the present study. At a first glance it would appear that, similar to signature calls being found in other mammals and birds (e.g., bottlenose dolphins: Janik & Slater, 1998; common marmosets: Jones et al., 1993; chiffchaffs: Naguib et al., 2001), the courtship songs of laboratory mice also contain individual signatures which vary between strains and could potentially serve for signalling identity and for promoting individual recognition by the female.

However, differently from the preliminary indications from Holy and Guo (2005), our hierarchical cluster analysis assessing the similarity across the whole song sequences revealed a remarkable individual variation in how similar courtship songs actually are within the same individual. We observed a gradient ranging from low to high individual consistency over time, and this finding could be generalised across the two different mouse strains. The concept of intra-individual variability, intended as a short-term, reversible behavioural variability of an individual in response to being repeatedly exposed to the same context (Nesselroade, 1991; Ram & Gerstorf, 2009; J. A. Stamps et al., 2012) may help explain our finding. Individuals can differ in how sensitive they are to different external stimuli (Aron & Aron, 1997; Dingemanse et al., 2010; J. Stamps & Groothuis, 2010) so that a context that is presented multiple times (i.e., repeated presentation of fresh female urine) may be perceived as the same or as a different one depending on the individual’s sensitivity to subtle changes (e.g., urinary cue coming from a different female in each test). Intra-individual variability can be regarded as differences in behavioural plasticity that are stable across individuals; such differences may be maintained by natural selection as they can contribute to the stability and persistence of a population in the face of a changeable environment (Dingemanse & Wolf, 2013). Thus, it is possible that different courtship strategies consisting of showing either low, medium or high phenotypic plasticity in song characteristics may be equally rewarded by bringing about different but complementary advantages for mating and reproduction. On the one hand, individual consistency of song sequences over time can be an advantage to ensure individual recognition and identification by a potential mate (Simmons et al., 2002); on the other hand, individual plasticity in song composition and sequence entropy may increase the overall complexity of the singing repertoire of an individual, and in turn be more attractive to a potential mate (Chabout et al., 2015; Leitão et al., 2006; Morisaka et al., 2008). This hypothesis should be empirically and systematically tested, for example by assessing whether one strategy is consistently favoured over the other by different female mice.

The identification within the song sequences of recurring, non-random motifs, which were previously identified as an integrant part of courtship songs in mice (Holy & Guo, 2005), highlighted again a large individual variation. The analysis of the syllabic structure of the motifs revealed that in both the strains studied, motifs had low complexity on average (i.e., simple structure with many repeated syllables) and showed similar levels of repetition within a song. These motif characteristics may be typical of the mouse species (see also similar examples in Holy & Guo, 2005). The high repetition of identical syllables and consequent simple structure of the motifs may imply that songs containing motifs may increase the chance that a song is heard in a “noisy” environment (Brumm & Slater, 2006).

### Conclusions

Taken together, not only genotype but also individuality can affect courtship songs in male mice. Our measures of length, composition and complexity of syllabic sequences contribute significantly to the general understanding of vocal communication, and could be used as tools to address ecologically relevant questions related to mate choice and reproductive success.

Our finding that certain syllabic features of an individual are overall consistent over time supports preliminary evidence that individuals can be identified based on syntactical characteristics of their songs. However, the considerable variation across individuals in how similar their song sequences are over time, suggests the presence of complementary courtship strategies related to different levels of behavioural plasticity. The adaptive value of this diversity in vocal communication is an area for future study.

## ACKNOWLEDGMENTS

The authors would like to thank the Ministry of Innovation, Science and Research of the state of North Rhine-Westphalia (MIWF) for supporting the implementation of a professorship for behavioural biology and animal welfare to SHR (Project: “Refinement of Animal Experiments”).

Furthermore, this project was supported by grants from the Deutsche Forschungsgemeinschaft (DFG, German Science Foundation) to SHR (281125614/GRK 2220, Project B6).

We are also very grateful to Prof. Franz Goller for his insightful and constructive feedback on a preliminary version of this manuscript.

## Appendix A1 Additional Methods

### Cage Furnishing and Enrichment

Cages of male mice were provided with wood chips (TierWohl Super, J. RETTENMAIER & SÖHNE GmbH, Rosenberg, Germany) as bedding material, a nestlet (BIOSCAPE GmbH, Castrop-Rauxel, Germany; Experiment 1) or a paper towel (Experiment 2) as nesting material and a transparent red plastic mouse house (Mouse House™, Tecniplast Deutschland GmbH, Hohenpeißenberg, Germany) and a wooden stick (approximately 1.5 cm × 1.5 cm × 10 cm) as environmental enrichment.

Cages of female mice contained the same items used for the males, except for the bedding material which consisted of wood shavings (females, Allspan, Höveler GmbH & Co.KG, Langenfeld, Germany), and the nesting material which always consisted of a paper towel.

### Vocalisations Recording Setup

All recordings of vocalisations were obtained in a dedicated testing room under red light conditions. Each male mouse was individually tested directly in its home cage, which was positioned in a noise cancelling box made of cardboard (l × b × h: 64 × 36 × 34 cm). The inside of the box was covered with corrugated acoustic foam (thickness: 2.5 – 3.5 cm) to improve the quality of the recordings. To allow an easy observation of the mouse inside its home cage, the front of the box was open. A high quality condenser microphone (Avisoft UltraSoundGate CM16/CMPA, Avisoft Bioacoustics, Germany) was placed inside the box over the centre of the home cage, pointing downwards and at a distance of 16 cm from the cage floor. The microphone was connected to a recording device (Avisoft UltraSoundGate 416Hb, Avisoft Bioacoustics, Germany) controlled by the software Avisoft-RECORDER (Version 4.2.27). Frequency range was set to 5 – 150 kHz, sampling rate to 300 kHz and resolution to 16 bit (Ferhat et al., 2016).

### Experimental Procedures

The experimental procedures below were carried out during the dark active phase, under red light conditions. The testing order of males was randomised within each batch and was kept the same across social encounter and test / control sessions. Whenever the experimenter touched a female mouse or its urine, gloves were changed immediately afterwards to avoid odour contamination within the testing room and across housing rooms.

### Social Encounter

One week before testing, each male experienced a 20 minute social encounter session with an unfamiliar female to induce the emission of courtship vocalisations by the male (Dizinno et al., 1978; Nyby et al., 1978; Sipos et al., 1992). The females used for these sessions were either 5-HTT +/− mice with a C57BL/6J genetic background (N = 24, PND 58 – 234; Experiment 1) or C57BL/6 mice (N = 16, PND 216; Experiment 2). A different female was randomly assigned to each male and in Experiment 2, where 16 females were used for 24 males, males from the same batch were always presented with females from different cages.

The social encounter took place in a Makrolon type III cage divided lengthways into two equal compartments by a perforated and transparent Plexiglas divider. This cage had wood chips as bedding and one water bottle for each compartment. First, one female was transported from the female housing room to the testing room and placed in one cage compartment, then the assigned male was transported in its home cage from the male housing room to the testing room and placed in the other compartment. The small holes in the divider (diameter: 5 mm) allowed for olfactory, auditory and limited physical contact between the two animals (similar to Musolf et al., 2010). The social encounter cage was then placed in the noise cancelling box and vocalisations were recorded to monitor the successful initial emission of courtship songs in response to female presence. The experimenter sat in front of the noise cancelling box to habituate the males to her presence in the testing room, and to check for any signs of severe stress (e.g., repeated escape attempts) by the animals, which however never occurred. After 20 minutes, both animals were returned to their housing rooms, and a new social encounter cage was prepared for the next mice pair.

### Test and Control Sessions

At the beginning of a test session, each male mouse was transported in its home cage to the testing room and the cage was placed in the noise cancelling box. The experimenter then left the room for 20 minutes to allow the animal to habituate to the novel environment. Another habituation period of 10 minutes followed in which the experimenter sat in front of the noise cancelling box. Then, fresh female urine was collected on a cotton swab in the female housing room, as described in the Urine Collection section. The cotton swab was immediately taken to the testing room and inserted through the grid of the cage lid onto one end of the home cage floor. Vocalisations were recorded for the five minutes following the presentation of the cotton swab. During this time, the experimenter recorded with a stopwatch the time that the animal spent sniffing the cotton swab. Sniffing time was measured to assess whether the animal was interested and therefore aware of the presence of the cotton swab in the cage. It was defined as having the snout in direct contact with the cotton swab, which also included carrying the swab around with the mouth. After the five minutes had passed, vocalisation recording was stopped and the animal was moved to a clean home cage. The cage lid was not replaced but was cleaned with ethanol to avoid contamination with female urinary cues. The animal was then taken back to the male housing room.

Differently from the test sessions, in the control sessions the animal was presented with a clean cotton swab without female urine. To account for the variable time the males had to wait during urine collection in the test sessions (Urine Collection section), in the control sessions the initial 30 minutes of habituation to testing room and experimenter were followed by a randomly varying waiting time (5 – 30 minutes) during which the experimenter was outside the testing room.

If the mouse did not sniff the cotton swab during the whole test or control session, it was assumed that the mouse may have not been aware of the swab presence in the cage (e.g., because it was resting in the nest), thus the session was repeated either at the end of the day or on the next day, so that the same animal was always tested at a similar time of day across testing sessions. This occurred three times in Experiment 1 (2 × C57BL/6J and 1 × B6D2F1 mice) and two times in Experiment 2 (2 × DBA/2 mice).

### Urine Collection

The females used for urine collection were 5-HTT +/+ mice with a C57BL/6J genetic background in Experiment 1 (N = 16, PND 65 – 221 at testing) or 5-HTT +/+ and 5-HTT +/− mice with a C57BL/6J genetic background in Experiment 2 (N = 21 and 14, respectively; PND 75 – 350 at testing). This was done to have a similar genetic background among female urine donors, and based on evidence that females’ genetic background does not have to match the male’s one for courtship songs to be induced (Holy & Guo, 2005; Sugimoto et al., 2011). It has been shown that male mice emit less vocalisations in response to cues from familiar females (Musolf et al., 2010) and that the oestrous stage of the cue female might influence vocalisation production as well (Barthelemy et al., 2004; Hanson & Hurley, 2012). Consequently, urinary cues were always obtained from unfamiliar, non-oestrous females. A list of the donor females was created before the start of the experiment and was followed in the same order throughout the study. Urine collection was performed in the female housing room. First, one female was taken out from its home cage and a vaginal smear was obtained with a clean plastic loop to determine the oestrous cycle stage, according to the methods from Byers and colleagues (2012). Briefly, the smear was suspended in a drop of water on a slide and vaginal cells were immediately investigated with a microscope (100x magnification under bright field illumination without staining). The predominant presence of cornified epithelial cells over leukocytes and nucleated epithelial cells indicated that the female was in the oestrus stage. If this was the case, the female was returned to its home cage and the next female on the list was collected for vaginal smear. If the female was in a stage different than oestrus, it was immediately placed in a clean Makrolon type II cage the floor of which was covered with a new sheet of aluminium foil for urine collection. In case the female had not urinated after three minutes, handling procedures such as tail handling, manual restraint by the nape and/or gentle stroking of the ventral area (Nyby et al., 1979) were performed to elicit urination. If urine could not be obtained within seven additional minutes, the female was returned to its home cage and the next female on the list was collected. As soon as the female urinated, 25 to 50 μl of urine were collected (Holy & Guo, 2005; Musolf et al., 2010) and transferred to a clean cotton swab by pipetting half the volume of urine on the centre of each side of the swab. The swab was then placed in a plastic container and transported to the testing room, so that no more than five minutes passed between urine sample collection and testing.

To prevent the risk that the emission of vocalisations within a strain could be affected by urine samples from a specific female, the priority was to collect urine samples from as many females as possible. Therefore, urine samples were taken from the same females only when all females from the list had donated one urine sample, or the ones that had not were in oestrous that day.

### Intra- and Inter- Rater Reliability

In Experiment 1, all syllables were counted by the same experimenter (S. Siestrup). Intrarater reliability was assessed by recounting the syllables of one session for every set of five sessions analysed (20 % of all recordings; n = 10). It was randomly chosen which recording out of the last five was reanalysed. The intraclass correlation coefficient (ICC) was calculated using a two-way mixed design assessing the absolute agreement of the average of the counts (Landers, 2015; Shrout & Fleiss, 1979). The ICC was computed for the total counts of syllables and the counts of each syllable type. Reliability was evaluated based on the ICC estimate and on the lower bound of the 95% confidence interval (Koo & Li, 2016). Intra-rater reliability was excellent for all the syllable types counted (all ICC_average_ > 0.994; all CI lower bounds > 0.981).

In Experiment 2, syllables were counted by two experimenters (S. Siestrup and M. Peng). Intra-rater reliability of the second experimenter (M. Peng) was assessed by recounting the syllables of one session for every set of eight sessions analysed (12 % of all recordings; n = 7). Inter-rater reliability was assessed on eight randomly chosen sessions that were analysed by both experimenters. Reliability was evaluated in the same way as in Experiment 1. Intrarater reliability was good to excellent (all ICC_average_ > 0.946; all CI lower bounds > 0.728). Inter-rater reliability was good to excellent for *Complex, Up-FM, Flat, Chevron, 1 Step, 2 Steps, 3 Steps* and total number of syllables (ICC_average_ > 0.962; CI lower bounds > 0.796) and was moderate to excellent for *Down-FM* and *Reverse Chevron* syllable types (ICC_average_ > 0.911; CI lower bounds > 0.512). *>3 Steps, Short* and *Other* syllable types had a good average intraclass correlation (ICC_average_ = 0.853, 0.811, 0.792; respectively) but relatively poor lower bound of the 95 % confidence interval (CI lower bounds = 0.305, 0.171, −0.074; respectively). Therefore, the syllable type *Other* was excluded from further analysis as it also occurred very rarely (less than 0.05 % of all emitted syllables), while the reliability of *>3 Steps* and *Short* syllable types was considered acceptable since the relative proportions of emitted syllables between BALB/c and DBA/2 remained similar between Experiment 1 and 2 (see Fig. 4 and appendix A3).

### Comparison of Control and Test Sessions

In Experiment 1, related-samples Wilcoxon signed rank tests were used to compare the total number of syllables produced during test and control sessions, and whether the difference between control and test sessions depended on the order of these two conditions. The same test was used to compare the time the animals spent sniffing the swab during test and control sessions. Significantly fewer syllables were emitted during control sessions (M = 127.5, IQR = 431.0) than during test sessions (M = 916.0, IQR = 792.0; z = 3.71, p < 0.001). When considering only the animals for which the control session preceded the test session, significantly more syllables were produced during the test (M = 787.5, IQR = 772.0) than during the control (M = 7.0, IQR = 95.0; z = −3.06, p = 0.002). Instead, when the test session preceded the control one, the number of emitted syllable during the test only tended to be greater than control (M = 1017.5, IQR = 1016.0 vs. M = 418.0, IQR = 729.0; z = −1.73, p = 0.08), indicating a carry-over effect from the test day to the control day. There was no significant difference between the time spent sniffing the swab during control (M = 195.0 s, IQR = 92.0 s) and test sessions (M = 186.0 s, IQR = 77.0 s; z = −0.36, p = 0.72), indicating that the animals showed similar interest in the cotton swab in either condition, and their vocalisations occurred in response to the urinary cue.

In Experiment 2, based on the results from Experiment 1, the control session always preceded the test sessions. The same comparisons as in Experiment 1 were made, but since the control session was followed by three test sessions, related-samples Friedman’s tests were used. BALB/c mice emitted fewer syllables in the control session compared to the test sessions overall, but also produced comparatively fewer syllables in the third test session (X^2^(3) = 26.10, p < 0.001; significant Bonferroni-corrected comparisons: Control < Time 1, Time 2; Time 2 > Time 3). DBA/2 mice also emitted fewer syllables in the control session than in the test sessions, and this difference was maintained across all test sessions (x^2^(3) = 20.70, p < 0.001; significant Bonferroni-corrected comparisons: Control < Time 1, Time 2, Time 3). In BALB/c mice, the time spent sniffing the cotton swab was overall greater in the control session than in the test sessions (x^2^(3) = 17.57, p = 0.001; significant Bonferroni-corrected comparisons: Control > Time 1, Time 3; as a trend: Control > Time 2), while in the DBA/2 mice no difference was found between control and test sessions 0<^2^(3) = 4.09, p = 0.25). Again, these results indicate that the time spent exploring the cotton swab did not affect the emission of vocalisations.

## Appendix A2 Sequence Entropy and Sequence Length (Experiment 1)

**Appendix A2.**
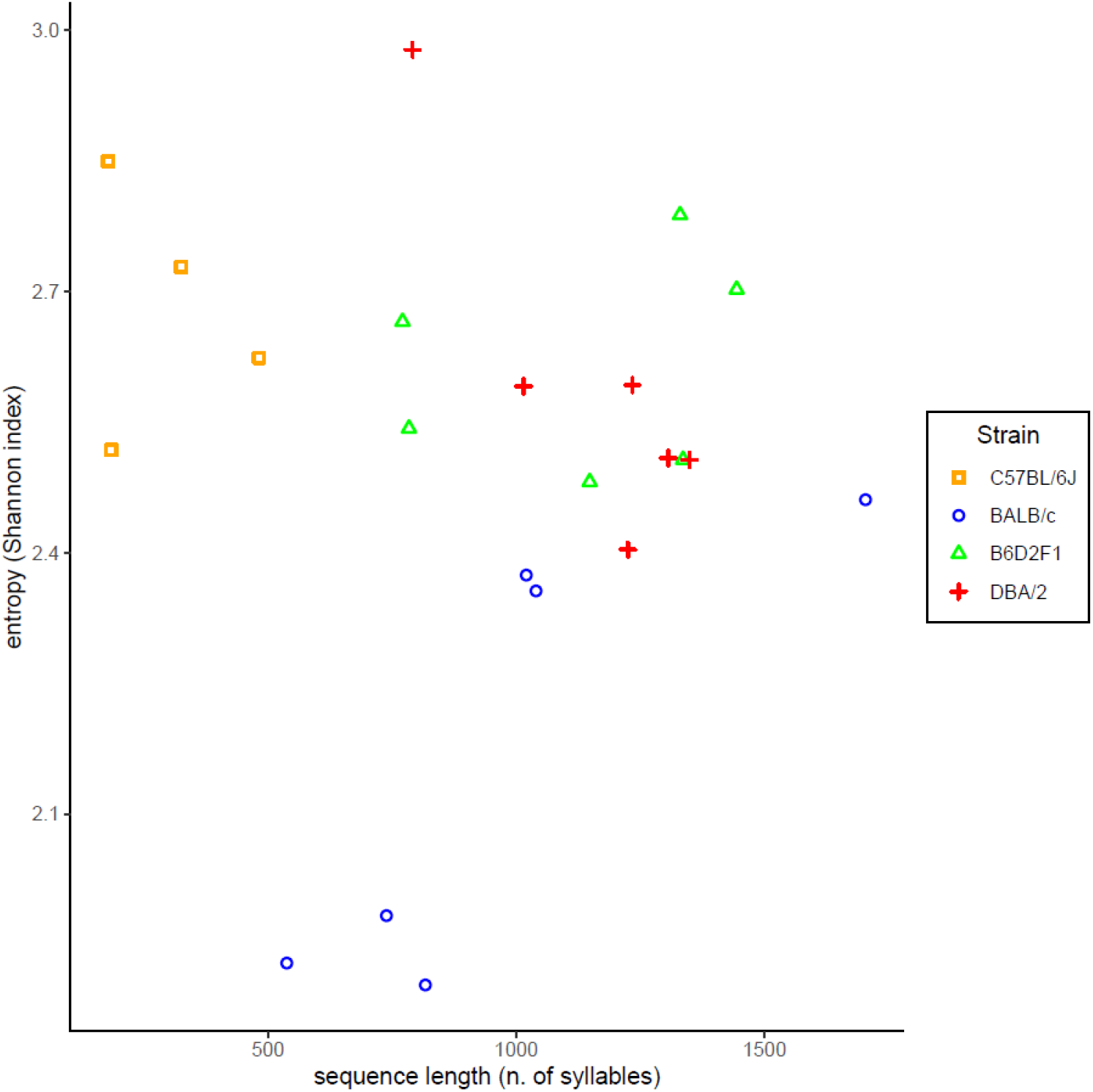
Scatter plot showing the relationship between sequence entropy and sequence length for each strain. Individuals (dots) from the same strain are assigned a dedicated symbol and colour. The sequence complexity of each strain appears to be differentially affected by these two measures.

## Appendix A3 Syllabic Composition (Experiment 2)

**Appendix A3.**
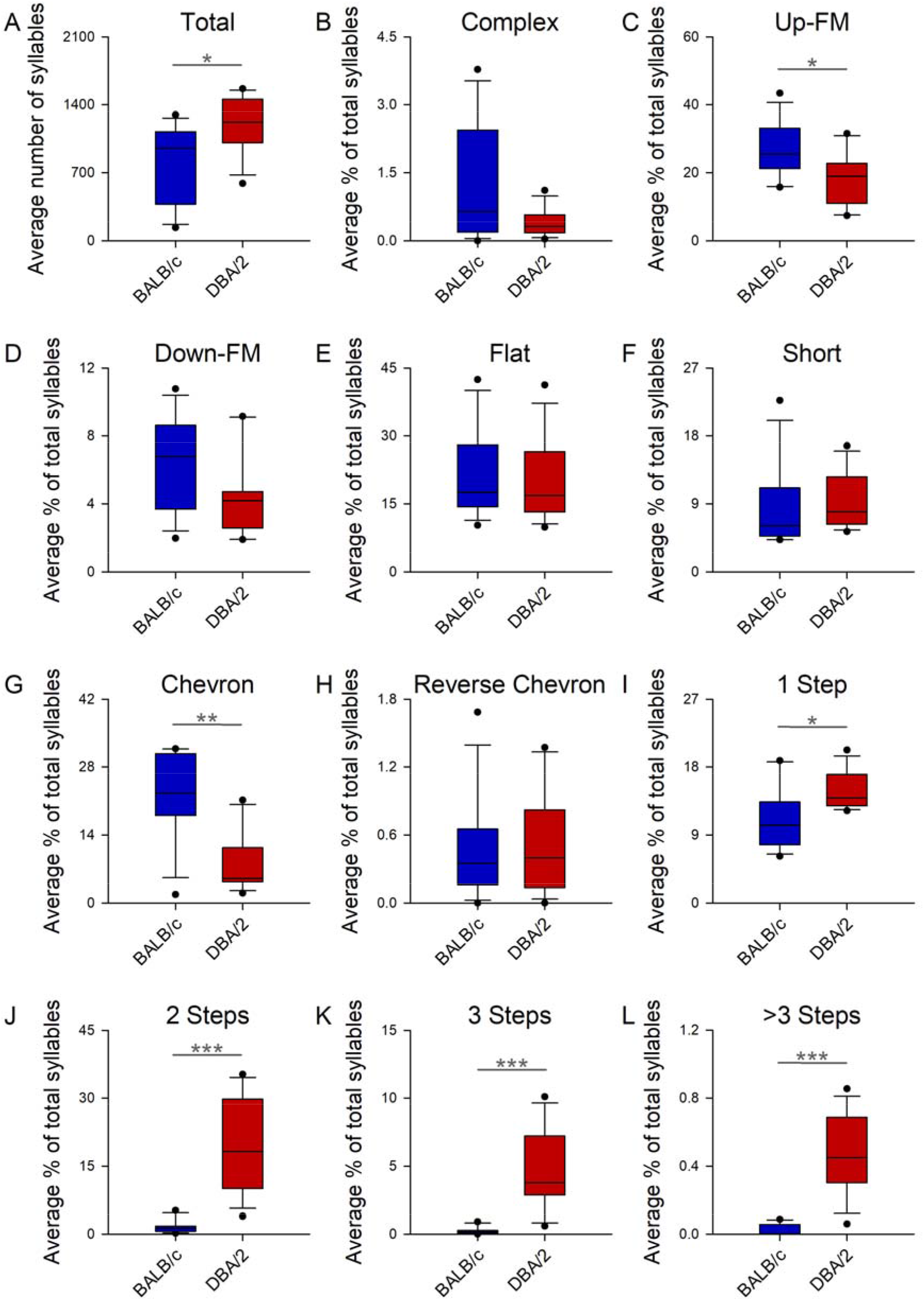
Box plots showing strain differences (BALB/c vs. DBA/2) in the total number of syllables and in the percentages of different syllable types emitted at testing (as average of the three test sessions). The boxes represent the middle 50% of the data, while the upper and lower whiskers include the middle 80% of the data; the upper and lower round dots indicate the 95th and 5th percentiles, respectively. T = p < 0.1, * = p < 0.05, ** = p < 0.01, *** = p < 0.001.

## Appendix A4 Temporal Consistency of Syllables

**Appendix A4.** Summary table showing the consistency over time of the total number of syllables emitted and of the four most frequent syllable types for each strain. Interpretation of the intraclass correlation (ICC) was based on the average ICC across the three test sessions and on the lower bound of the 95 % confidence interval; lower bounds below 0.5 indicated poor consistency (Koo & Li 2016). Lower bounds of the confidence interval which were above the 0.5 threshold are shown in bold.

| Strain | Syllable Type | Average ICC | 95 % Confidence Interval |  | ICC Interpretation* |
| --- | --- | --- | --- | --- | --- |
|  |  |  | Lower Bound | Upper Bound |  |
| BALB/c | *Total Syllables* | 0.859 | **0.626** | 0.956 | Moderate/Excellent |
|  | *Up-FM* | 0.734 | 0.220 | 0.928 | Poor |
|  | *Chevron* | 0.341 | -0.931 | 0.822 | Poor |
|  | *Flat* | 0.828 | 0.495 | 0.953 | Poor |
|  | *1 Step* | 0.415 | -0.714 | 0.842 | Poor |
| DBA/2 | *Total Syllables* | 0.624 | 0.006 | 0.883 | Poor |
|  | *2 Steps* | 0.823 | **0.531** | 0.945 | Moderate/Excellent |
|  | *Up-FM* | 0.915 | **0.775** | 0.973 | Good/Excellent |
|  | *Flat* | 0.653 | 0.082 | 0.891 | Poor |
|  | *1 Step* | 0.010 | -1.619 | 0.690 | Poor |

\*Based on Koo & Li (2016).

## Appendix A5 Examples of High or Low Temporal Consistency in Syllabic Composition

**Appendix A5.**
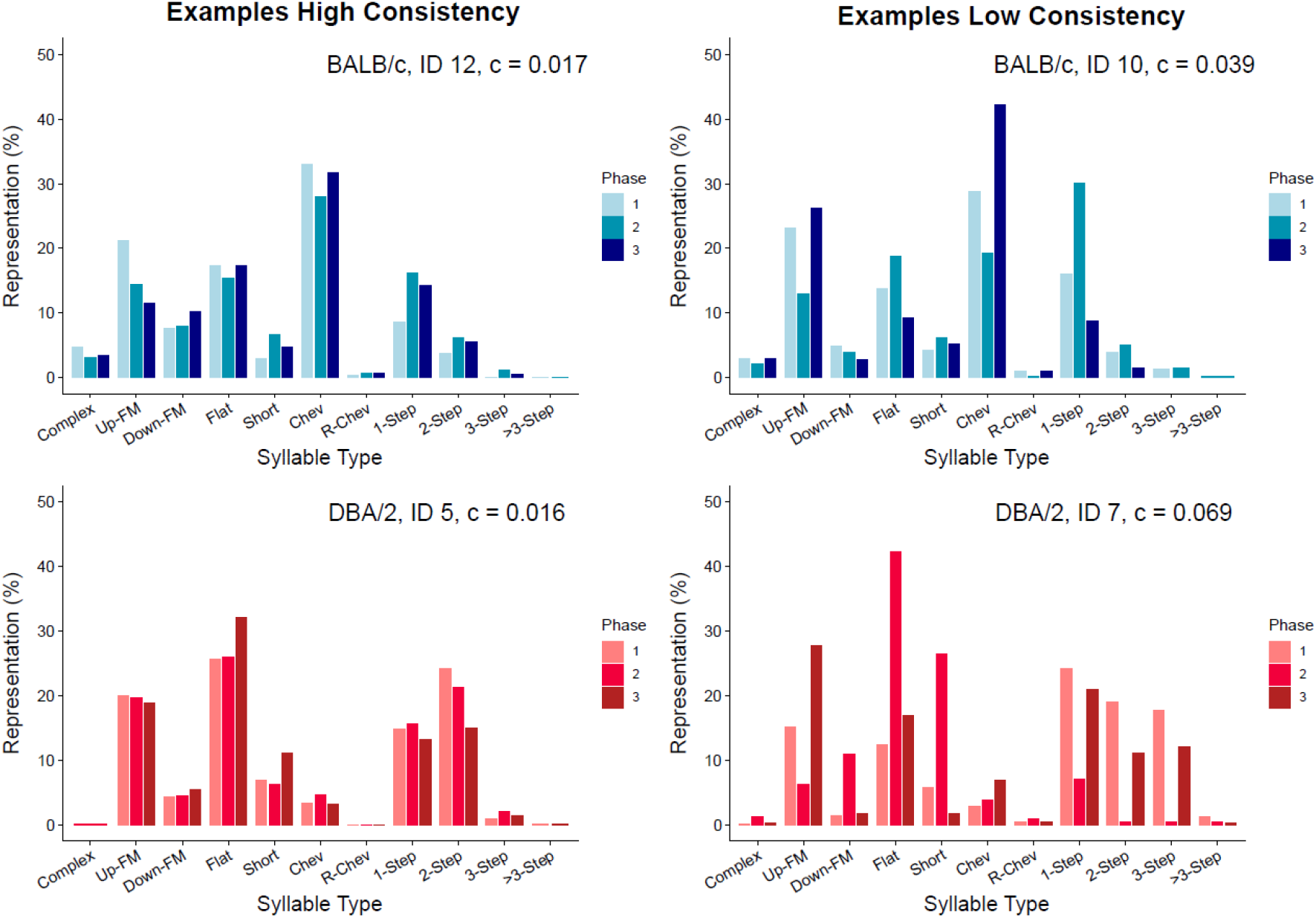
Examples of individual mice (two per strain) showing high or low temporal consistency in syllabic composition of courtship songs. For each individual, the consistency score “c” was determined by first calculating the standard deviation of the percentages of each syllable type across the three test sessions, and then averaging these standard deviations across all syllable types. The smaller the c score, the more consistent is the syllabic composition across test sessions.

